# Cellular thermal shift assay of subcellular isolates for evaluating drug-membrane target interactions

**DOI:** 10.64898/2026.02.03.703651

**Authors:** Sahiba K. Dogra, Varunya Kattunga, Shona A. Mookerjee, Anand Rane, Manish Chamoli, Julie K. Andersen

**Author notes:** Corresponding Author: Dr. Julie K Andersen, Buck Institute for Research on Aging, 8001 Redwood Blvd, Novato, CA 94945, United States of America.

## Abstract

The cellular thermal shift assay (CETSA) is an invaluable tool for target identification and validation in early drug discovery efforts. It relies on thermal melting curves to indicate drug binding and is typically performed in whole cells, cell lysates, or purified protein as validation of direct interaction. However, these approaches can result in disruption of the structural integrity of membrane proteins, hindering downstream analysis and drug-target engagement. Here, we describe the first application of CETSA in isolated mitochondria and show the effects of this approach on the analysis of the compound UK5099 and its known binding target, the mitochondrial pyruvate carrier (MPC), a mitochondrial inner membrane-localized protein complex. Our analysis supports a model in which the MPC must remain structurally intact for UK5099 binding. We demonstrate that the binding of UK5099 to the MPC is disrupted in whole cells and cell lysates, whereas isolating mitochondria maintains the binding interaction between drug and target observable using CETSA. These data suggest that isolating membrane-bound organelles through subcellular CETSA stabilizes membrane-bound proteins in their native conformation, allowing the identification of membrane-localized drug binding targets that might otherwise be missed.

## Introduction

A clinically successful drug must specifically modulate its target to produce a beneficial therapeutic effect while minimizing any off-target effects. Methods to validate drug-target interactions include surface plasmon resonance (SPR), nuclear magnetic resonance (NMR), and X-ray crystallography. However, these techniques often use purified (or crystallized) protein, a critical limitation for the analysis of membrane proteins and complexes. Membrane proteins require a lipid environment to maintain their native structure which can be disrupted by solubilization via detergents and is difficult to artificially configure outside of the cell. Traditional drug-target validation methods may therefore constrain membrane-bound proteins to a non-physiological state that may not accurately represent that present during drug binding *in vivo*, impeding drug discovery [1]. Methods for assessing drug-target interactions that preserve membrane proteins in a more physiologically-relevant state are therefore urgently needed.

Unlike NMR, SPR, and X-ray crystallography, the cellular thermal shift assay (CETSA) is a label-free method that can be performed in whole cells and cell lysates in addition to purified protein, providing a comprehensive assessment of drug binding in various contexts [2]. CETSA utilizes temperature-dependent protein denaturation to assess the effects of drug binding on protein thermal stability. Drug interaction with a protein target can either stabilize or disrupt the target structure. The change in the structural stability can in turn affect protein susceptibility to thermal denaturation. Since denatured proteins precipitate into the insoluble fraction during heat treatment, measuring the amount of protein present in the soluble fraction across a given temperature range yields that protein’s thermal curve and melting temperature (T_m_) - the temperature at which 50% of the protein present in the soluble fraction at physiological temperature denatures. The drug-target interaction is then visualized as a shift in the T_m_ [3].

CETSA has become ubiquitous in early drug discovery pipelines to identify and validate primary drug-protein targets [3]. However, its use for the analysis of membrane proteins as drug targets is limited by losses in structural integrity during the solubilization process due to removal from their native membrane environment [4]. Membrane proteins such as G-protein coupled receptors or membrane transport proteins are positioned at key therapeutic signaling interfaces, increasing the importance of a drug-target validation method for such targets [1].

In a standard CETSA assay, cultured cells are exposed to the compound of interest. Next, the cells are incubated at a range of temperatures and lysed before the soluble fraction is isolated. There are several reasons that CETSA has been a relatively poor approach to analyzing membrane-bound proteins. First, when CETSA is performed in cultured cells, membrane proteins typically denature and precipitate into the insoluble fraction in the cell lysis step after thermal exposure, preventing analysis of the soluble fraction in the context of thermal stability. This issue been recently addressed by Kawatkar et al., who reported that use of low detergent concentration can preserve membrane protein structure during downstream processing of samples following whole cell CETSA without affecting the thermal stability of membrane proteins. This innovation allows for assessment of drug-target interactions for membrane proteins in the context of a whole cell environment [4]. However, the interpretation of whole cell CETSA can be compromised when the drug affects the thermal shift of a protein indirectly via upstream regulation of transcription, translation, and/or post-translational modifications or when cellular pathways/structures affect drug uptake or binding. Hence direct drug-protein engagement via CETSA is usually performed in the context of cell lysates or the purified protein target [3]. Removing membrane proteins from the native membrane can disrupt the native conformation of membrane proteins including those located in intracellular organelles [1]. The drug is therefore not able to bind to disrupted membrane proteins in cellular lysates or in their purified states, producing false negative results [4].

To overcome these confounds, we applied CETSA to subcellular isolates in this case in isolated mitochondria (Table 1). Subcellular isolation is technically straightforward, providing an easy-to-adopt methodology for expanding target validation efforts to sensitive protein targets. As a proof-of-principle, we chose to detect the well-established binding of the drug UK5099 to its known target, the mitochondrial pyruvate carrier (MPC) complex. UK5099 is used in clinical trials to stress test human mitochondrial samples and evaluate mitochondrial function [5]. UK5099 has been shown to bind to the MPC through differential scanning fluorimetry and recently has been further validated through detailed structural analysis through cryo-electron microscopy [6, 7]. The MPC consists of a heterodimer formed by MPC1 and MPC2, two transmembrane proteins spanning the mitochondrial inner membrane (MIM). UK5099 binds to the complex at a site in the pocket of the MPC heterodimer, requiring that the complex remain intact for binding to occur [6]. Our data demonstrate that while drug-induced heat stabilization of the MPC is not detectable in CETSA performed on whole cells or cellular lysates, CETSA performed in isolated mitochondrial fractions demonstrated the expected drug-induced thermal shift of the native MPC heterodimer. These findings demonstrate that performing CETSA on mitochondrial isolates is a viable means of overcoming limitations of CETSA on whole cell and cell lysates. This method can likely be extended to other organellar isolates such as the endoplasmic reticulum whose membrane proteins are critical for cell signaling and organellar function in disease states [1].

**Table 1:** Comparison of CETSA application in different biological preparations. Whole cell CETSA, whole cell lysate CETSA, and purified protein CETSA present limitations in the context of membrane-bound proteins that can be directly addressed via subcellular CETSA.

| Method | Pros | Cons |
| --- | --- | --- |
| Whole cell CETSA | • Most biologically relevant context | • Complexity of whole cell environment can yield artifacts<br>• Membrane proteins are difficult to preserve under experimental conditions |
| Cell Lysis CETSA | • Removes majority of confounding cellular environment effect on protein's thermal stability | • Membrane proteins are disrupted by lysis process<br>• Less biologically relevant context |
| Purified Protein CETSA | • Only factor affecting thermal stability is ligand binding | • Expensive<br>• Time-consuming<br>• Can be technically challenging for membrane proteins<br>• Least biologically relevant context |
| **NEW:** Subcellular isolate CETSA | • Maintains structure of mitochondrial membrane<br>• Maintains intact mitochondria, preserving biologically relevant context | • Requires higher cell quantity for isolation and sample prep |

## Methods

### 2.1 Cell lines and cell culture

HEK293FT cells (ThermoFisher Ref R70007) were cultured in Dulbecco’s Modified Medium (DMEM, Corning Ref 10-013-CV) supplemented with 10% fetal bovine serum (FBS, Corning Ref 35-010-CV) and 10% penicillin-streptomycin (pen-strep, Gibco Ref 15140122) at 37°C in a 95% humidity environment containing 5% CO_2_. Cells were allowed to reach near confluency (∼90%) in a T-75 (Genesee Scientific 25-209) flask before being transferred to a second T-175 flask (Genesee Scientific 25-211). Upon reaching near confluency in the second flask, the cells were split between four T-175 flasks.

### 2.2 Mitochondrial enrichment

Approximately 170-200 million HEK293FT cells were trypsinized for 2 minutes with 0.25% trypsin (Corning Ref 25-053-Cl) and quenched with equal amounts of culture medium. Cells were centrifuged at 300 x *g* for 3 minutes. The cell pellet was resuspended in Dulbecco’s phosphate-buffered saline (VWR 02-0119-0500) and cells were centrifuged again at 300 x *g* for 3 minutes. The resulting cell pellet was resuspended in pre-chilled mitochondrial isolation buffer [320 mM sucrose (Avantor Ref 4097-06), 1 mM EGTA (Millipore Sigma (CAS 67-42-5), 5 mM TES (Millipore Sigma (7365-44-8), pH 7.2] at a volume of 5 times the volume of the pellet. To minimize protease activity, the following protocol was performed on ice. The cell suspension was transferred into a pre-chilled Dounce homogenizer and homogenized with 30 to 50 strokes. The homogenate was transferred into a 15 mL tube and centrifuged at 800 x *g* for 10 minutes at 4°C. The supernatant was saved. The pellet was resuspended in mitochondrial isolation buffer at a volume 5 times the size of the pellet before the suspension was transferred to a Dounce homogenizer and homogenized using at least 30 strokes. The homogenate was centrifuged at 800 x *g* for 10 min at 4 °C. The supernatant was combined with the supernatant from the first spin and centrifuged for 12,000 x *g* for 15 minutes at 4°C [8]. The final supernatant was saved for analysis of mitochondrial membrane integrity by western blot to assess whether mitochondrial membrane proteins were present in the supernatant. The pellet containing the mitochondria was resuspended in mitochondrial isolation buffer at 1x pellet volume for use in CETSA analysis.

### 2.3 CETSA

#### 2.3.1 CETSA on isolated mitochondria

Protein concentration of mitochondrial isolates was determined by Bradford assay. Approximately 10% of the mitochondria from a given isolation was transferred to a new tube with equal volume of cell lysis buffer [150 mM NaCl (Millipore Sigma CAS 7647-14-5), 1% NP-40 (Millipore Sigma CAS 9002-93-1), 50 mM Tris-HCl (Millipore Sigma CAS 1185-53-1), pH 8.0, protease inhibitor (Millipore Sigma Ref 11-836-170-001), and phosphatase inhibitor (Millipore Sigma Ref 04-906-837-001)]. The sample was thoroughly triturated before quantifying the protein via Bradford Assay. BSA standards (BioRad 5000206) were prepared in triplicate from 0 to 4 mg/mL in a 96 well plate. Bradford reagent (Bio-Rad Ref 5000006) was prepared in a 1:5 dilution in MilliQ water before vacuum filtration (Genesee Scientific 25-231). Then, 200 µL of Bradford reagent was added to each well. Using a spectrophotometer, the plate was read at 595 nm. The total amount of protein in the mitochondrial pellet was then estimated from the 10% lysed extract. The remaining lysed extract was used for downstream Western blot analysis of mitochondrial membrane integrity and purity. Aliquots of 100 µg of mitochondrial protein per condition were distributed among 0.2 mL tubes. Traditional loading controls like ß-actin are not suitable for CETSA since they also denature at higher temperatures. The blots are instead normalized by loading an equal amount of protein per sample.

UK5099 (Cayman Chemical Company Ref 16980) was added at a final concentration of 10 µM in a final volume of 50 µL per condition. Upon adding UK5099 or DMSO vehicle (Millipore Sigma Ref D2650) alone, the samples were incubated on a platform shaker at 1000 rpm for 90 minutes. After 90 minutes, a PCR cycler (Bio-Rad C1000 Touch Thermal Cycler) was used to heat the samples for 3 minutes across 9 temperatures (37, 42, 44.1, 47.3, 51.4, 56.9, 61.1, 64, 66°C). Samples were then cooled at room temperature for 3 minutes [3].

Cell lysis buffer was added to each sample for a final concentration of 0.2% NP40 and 0.2% protease and phosphatase inhibitors in a final volume of 50 µL. The samples were mixed well before being lysed in liquid nitrogen for a total of three freeze thaw cycles. The samples were then centrifuged at 20,000 x *g* for 20 minutes. Supernatant containing soluble proteins were moved to a new tube for downstream western blot analysis [4].

#### 2.3.2 CETSA on mitochondrial lysates

Aliquots of 100 µg mitochondrial protein were lysed with cell lysis buffer and liquid nitrogen before compound incubation as described above. The soluble fraction was isolated as described above.

#### 2.3.3 CETSA on whole cells

Whole cells were treated with 10 µM UK5099 for 90 minutes. The cells were harvested at 75 million cell mL^-1^ in PBS with protease and phosphatase inhibitors at a concentration of 1% [4]. Cells were then incubated and lysed as above. The soluble fraction was isolated as described above.

#### 2.3.4 CETSA on cellular lysates

Cells were isolated at 62.5 million cells mL^-1^ in PBS with protease and phosphatase inhibitors. The cells were lysed in cell lysis buffer and liquid nitrogen as described above. Then, compound or vehicle control was added for 90 minutes in a final volume of 50 µL. Incubation and soluble fraction isolation proceeded as described above.

### 2.4 Western blot analysis

Sodium dodecyl sulfate-polyacrylamide gel electrophoresis (SDS-PAGE) was used both for CETSA analyses and for validation of mitochondrial enrichment. A 12 % gel (Invitrogen Ref NP0341BOX) was run in MOPS buffer (Invitrogen Ref NP0001) at 80-100 V for an hour and a half. The gel was transferred using 10x Transfer buffer [1.92 M Glycine (Sigma Ref 56-40-6), 0.25 M Tris (Sigma Ref 77-86-1)] and a methanol-activated (Millipore Sigma Ref 179337) PVDF membrane (Millipore Sigma Ref ISEQ00010). The transfer was run at 250 mA for 90 minutes. The membrane was blocked in 2% BSA for two days before being probed with 1:500 MPC1 antibody (Cell Signaling Ref 14462) [9], 1:2000 MPC2 antibody (Cell Signaling Ref 46141) [10], 1:2000 Porin VDAC antibody (Calbio Ref 529534), or 1:2000 ß-actin antibody (Cell Signaling Ref 3700) overnight. The secondary goat anti-rabbit IgG antibody for MPC1 and MPC2 was diluted 1:10,000 (Millipore Sigma Ref AP132P) and the secondary goat anti-mouse IgG antibody for the other proteins were diluted 1:2000 (Millipore Sigma Ref AP127P). The membranes were visualized using chemiluminescent HRP substrate with either normal ECL (Cytiva Ref RPN2106) or enhanced ECL (Millipore Sigma WBKLS0500) and the Bio-Rad ChemiDoc Imaging System.

### 2.5 Western blot quantification and statistical analysis

Image Lab BioRad software (version 6.1) was used to quantify the Western blot images. The rectangle ROI tool and the analysis table were used to obtain the mean intensities. The area of the analyzed field was held constant across measurements within a blot. All measurements were normalized to the band intensity at 37°C, such that protein abundance in the 37°C condition corresponds to 100%; 0% was defined as the temperature at which all the protein was insolubilized. Normalized values were fit to the Boltzmann Sigmoid Curve using GraphPad PRISM (version 7) [3]. Statistical significance between treatments was evaluated using an extra sum of squares F-test if the melting curves met the parameters to be fitted to a Boltzmann Sigmoid equation or a t-test assuming unequal variance if one or both curves being compared did not meet the parameters for curve fitting. Technical replicates are defined as n and biological replicates are defined as N.

## Results and Discussion

### Design and synthesis of mitochondrial isolation and CETSA

We designed a novel subcellular CETSA using mitochondria as a proof-of-principle to overcome limitations of the CETSA approach in whole cell and cell lysates (Figure 1). Our primary goal was to preserve the analysis of the drug-target interaction potentially lost in whole cell and cell lysate samples. We therefore decided to interrogate whether enrichment of structurally intact mitochondrial isolates was sufficient for investigating drug-target interactions via CETSA. Crude mitochondrial isolation also allows for larger extractions and a greater sample preparation, allowing for numerous CETSA samples to be processed in parallel. With this approach, negative controls, positive controls, and unknown drug samples can all be processed using the same mitochondrial preparation, thereby minimizing variability. To measure whether mitochondria remain structurally intact following enrichment, we compared the presence of mitochondrial protein in the pellet to the supernatant in the final centrifugation step. If the mitochondrial membranes are lysed/denatured during the purification process, significant levels of mitochondrial membrane proteins would be expected to be present in the supernatant as well as the mitochondrial fraction. We did not see VDAC or MPC1 in the supernatant of our crude mitochondrial isolate. ß-actin was present in both the supernatant and mitochondrial pellet, demonstrating that samples represent enrichment rather than full purification of the mitochondrial fraction (Supplementary Figure 1).

**Figure 1:**
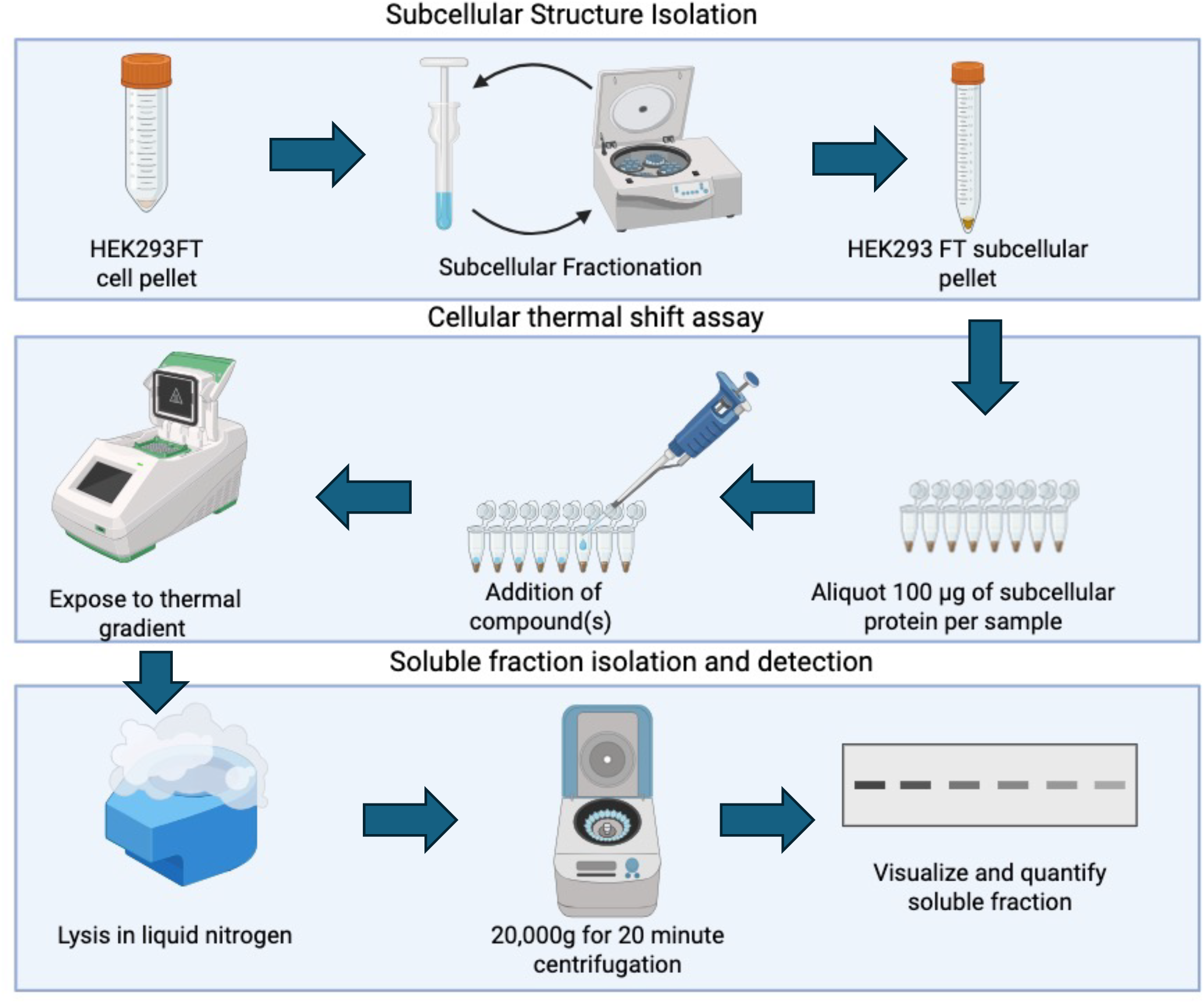
Schematic of performance of CETSA on subcellular isolate. The procedure is divided into three parts: subcellular structure enrichment, drug treatment and CETSA, and finally isolation and analysis of the soluble fraction.

Considerations for use of the protocol include the amount of mitochondrial protein needed per sample depending on the protein abundance, incubation time, and the thermal range. For a target in the inner membrane such as the MPC, the drug may require a longer incubation time than required for a protein localized in the outer mitochondrial membrane. Each protein target will have a different thermal range as the final temperature at which all the protein becomes insolubilized depends on the protein’s innate thermal stability.

### 3.1 CETSA in both whole cell and cellular lysate environments disrupts UK5099-MPC1 binding interaction

To assess the impact of MPC1/MPC2 drug binding in the context of cellular lysates versus crude mitochondrial extracts, we used the well-established inhibitor of MPC activity UK5099 in our CETSA analyses. UK5099 interacts with the binding pocket located between the two MPC monomers making up the heterodimer, requiring maintenance of structural integrity of the entire protein complex [6]. These characteristics make the UK5099-MPC interaction highly challenging to examine in the context of whole cell or cellular lysate CETSA. No statistically significant thermal shift of the MPC1 melting curve in response to UK5099 treatment was observed in whole cells (T_m_ of MPC1 = 61.4 ± 2.9 °C in drug-free controls versus 60.9 ± 3.1 °C in UK5099 treated samples). The higher T_m_ of MPC1 indicates that the structure of the protein itself is preserved, but the absence of a significant thermal shift in the presence of UK5099 indicates that the drug-target interaction is not detectable (Figure 2a, 2b). Since it is known that UK5099 binds to the MPC, the downstream processing including the cell harvesting and lysis of the membrane proteins likely masked the thermal shift that occurred during target engagement.

**Figure 2:**
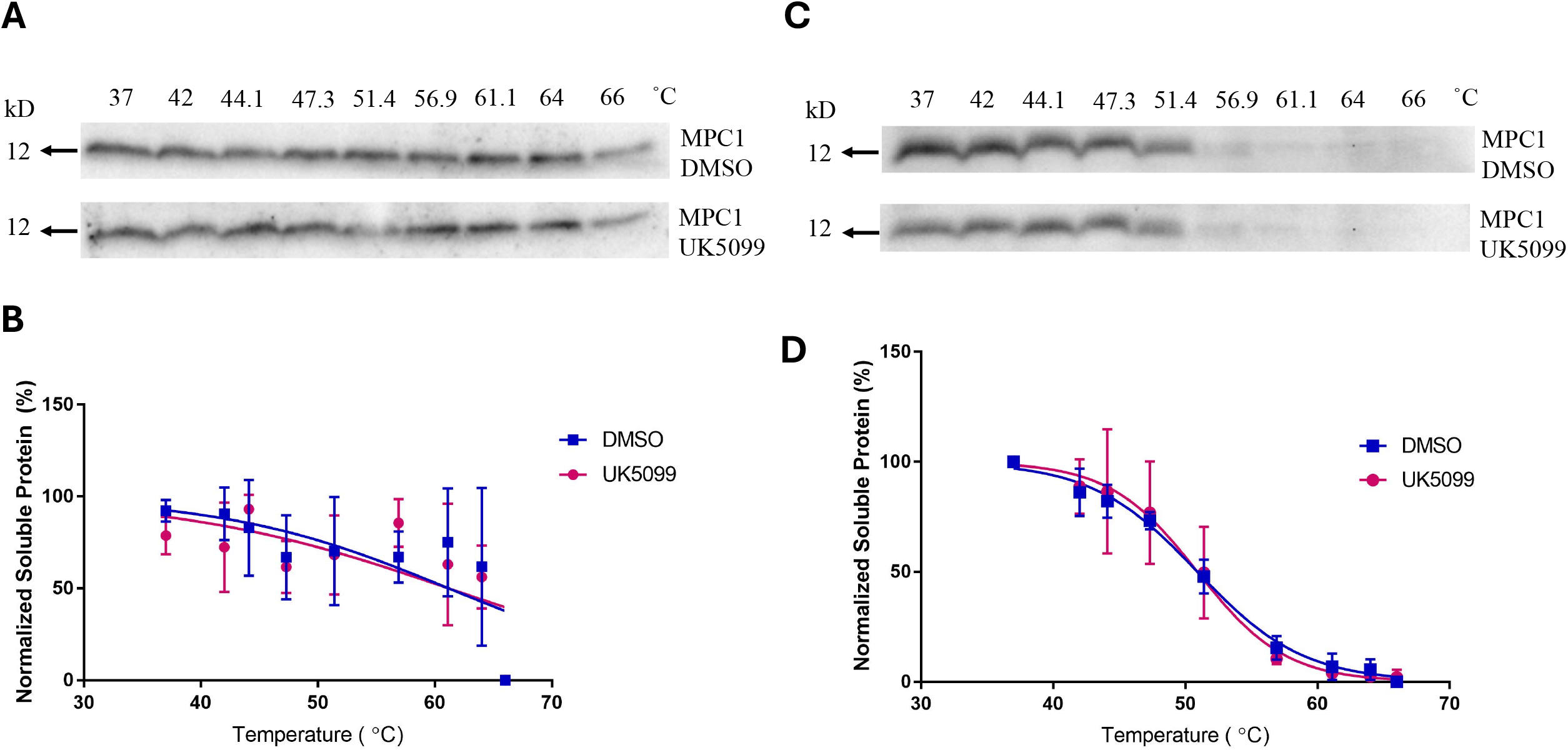
No detection of UK5099:MPC1 interaction in CETSA performed on whole cells or whole cell lysate. (A) Western blot image of MPC1 at 12 kD in the soluble fraction from 37 to 66°C of whole cells treated with vehicle control and UK5099. (B) The western blot was quantified using Image Lab.Relative intensities were used to quantify protein abundance normalized to 37°C. Curves fitted to the Boltzmann Sigmoid equation and demonstrate no statistically significant difference between the melting temperature of MPC1 in the control and UK5099-treated conditions (n = 1, N = 3, extra sum of squares F test, p = 0.91). (C) Western blot image of MPC at 12 kD in the soluble fraction of whole cell lysate treated with vehicle control and UK5099. (D) Quantified protein abundance in the soluble fraction fitted to the Boltzmann Sigmoid equation demonstrate no significant difference in the melting temperatures between control and UK5099-treated conditions (n = 1, N = 3, extra sum of squares F test, *p* = 0.82). Blot images are representative of means obtained from independent replicates.

Similarly, in the cellular lysate context, no statistically significant thermal shift of the MPC1 melting curve in response to UK5099 treatment was observed (T_m_ of MPC1 = 50.8 ± 0.3 °C in drug-free controls versus 50.9 ± 0.7 °C in UK5099 treated samples). The lower T_m_ in the cellular lysate context indicates that the structure of MPC1 itself was compromised by the lysis process utilizing detergent and freeze thaw cycles (Figure 2c, 2d). The false negative result in both traditional contexts misrepresents the strong binding of UK5099 to MPC indicated by an IC50 value of approximately 52.6 nM [7] and indicates that binding has been masked. This is likely due to the effects of lysis on the native conformation of the protein complex, resulting in altered protein solubility and disruption of the binding pocket and heterodimer formation.

### 3.2 Detection of thermal shift in MPC1 and MPC2 following addition of UK5099 is restored in mitochondrially-enriched samples

Next, we assessed UK5099-MPC drug binding in the context of CETSA on crude mitochondrial extracts. Establishing a robust thermal curve and aggregation temperature for a protein target in control conditions is required for subsequent CETSA analysis. Therefore, we first determined the control thermal curve and aggregation temperature of both MPC1 and MPC2 in the mitochondrial isolates (Figure 3a, 3c). The melting curve of MPC1 and MPC2 fit the Boltzmann Sigmoid curve for thermal denaturation and the T_m_ of MPC1 and MPC2 are 60.5 ± 0.7 °C and 57.1 ± 1.3 °C respectively. We observed that the T_m_ of MPC1 in the crude mitochondrial isolates (T_m_ = 60.5 ± 0.7 °C) is approximately the same as the T_m_ in whole cells (T_m_ = 61.4 °C ± 2.9) but 10°C higher than the aggregation temperature in cellular lysates (T_m_ = 50.8 ± 0.3 °C). The similarity in T_m_ in the whole cell and mitochondrial isolate conditions suggests that the MPC1 conformation is physiologically relevant and structurally maintained. The difference in T_m_ in the cellular lysate context suggests that the MPC is better maintained in its native conformation in the mitochondrial isolate context.

**Figure 3:**
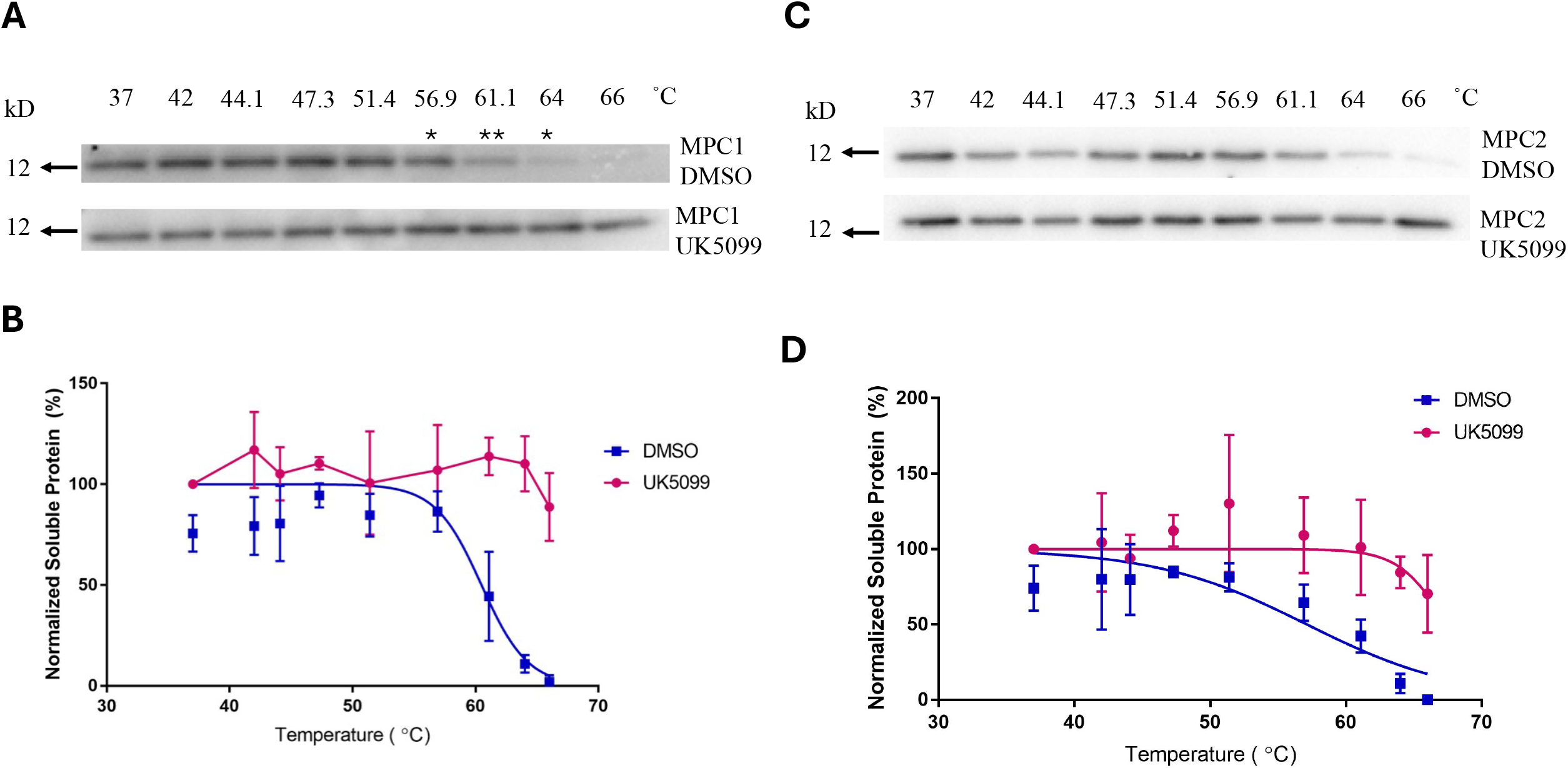
Detection of UK5099:MPC1 interaction in CETSA performed on crude mitochondrial isolate. (A) Western blot image of MPC1 in the soluble fraction between the temperatures of 37 and 66°C in DMSO and UK5099-treated conditions. (B) The western blot was quantified as described above. The DMSO samples were fitted to a Boltzmann Sigmoid curve and compared to the UK5099 samples (n = 1-2, N = 3, paired t-test assuming unequal variance, * = p < 0.05, ** = p < 0.01). (C) Western blot image of MPC2 in the soluble fraction between the temperatures of 37 and 66°C in DMSO and UK5099-treated conditions. (D) The western blots were quantified as above and the DMSO samples were fitted to a Boltzmann Sigmoid curve and compared to the UK5099 samples (n = 1-2, N = 3, extra sum of squares F-test, p < 0.05). Blot images are representative of means obtained from independent replicates.

To assess the use of mitochondrial extracts to overcome the limitations of whole cell and cellular lysate CETSA, we next examined this in the context of UK5099 treatment. We observed a significant thermal stabilization of MPC1 and MPC2 demonstrating that roughly 90% and 70% of these two proteins, respectively, were stabilized in the soluble fraction at 66°C, the highest temperature examined. The stabilization reflects the established strong binding of UK5099 to MPC [7]. Given the strong stabilization of MPC1, the data did not meet the criteria for fitting to the Boltzmann Sigmoid curve. Instead, we compared the last several temperatures of MPC1 using a t-test to establish statistical significance of the stabilization observed at higher temperatures. For MPC2, we observed a significant 10°C shift in the target’s thermal stability (T_m_ of MPC2 = 57.1 ± 1.3 °C in drug-free controls versus T_m_ of MPC2 = 67.4 ± 2.3 °C in UK5099 treated samples). As opposed to the lack of thermal shift observed in the context of whole cell or cell lysates, thermal stabilization of both MPC1 and MPC2 emulates what is expected to occur as a consequence of the established interaction of UK5099 in the binding pocket between the two monomers (Figure 3b, 3d). Taken in total, our data demonstrates that isolation of mitochondria allows for a more sensitive analysis of drug-protein interactions in the membrane which can be applied to other subcellular compartments.

### 3.3 Lysis of the mitochondria results in loss of thermal MPC1 shift in the presence of UK5099 observed in mitochondrially enriched samples

To confirm that drug-target interaction requires mitochondrial integrity, we performed the same subcellular CETSA but lysed the mitochondria prior to rather than after drug treatment (Figure 4a). The UK5099-MPC interaction observed following CETSA in mitochondrially-enriched fractions was lost following lysis. No statistically significant differences in the thermal curve were observed between DMSO control (T_m_ = 54.6 ± 1.4 °C) and UK5099 (T_m_ = 53.94 ± 1.7 °C) in lysed mitochondrial samples. The lack of a thermal shift in UK5099-treated lysed mitochondria indicates that lysis of the mitochondrial membrane is responsible for the false negative result in CETSA performed in lysed samples (Figure 4b, 4c). The thermal stabilization observed in isolated mitochondria is therefore due to preservation of membrane integrity rather than an artifact of compound enrichment or sample preparation. We concluded that subcellular isolation enhances thermal shifts by maintaining the integrity of the target protein during drug-target engagement and downstream processing.

**Figure 4:**
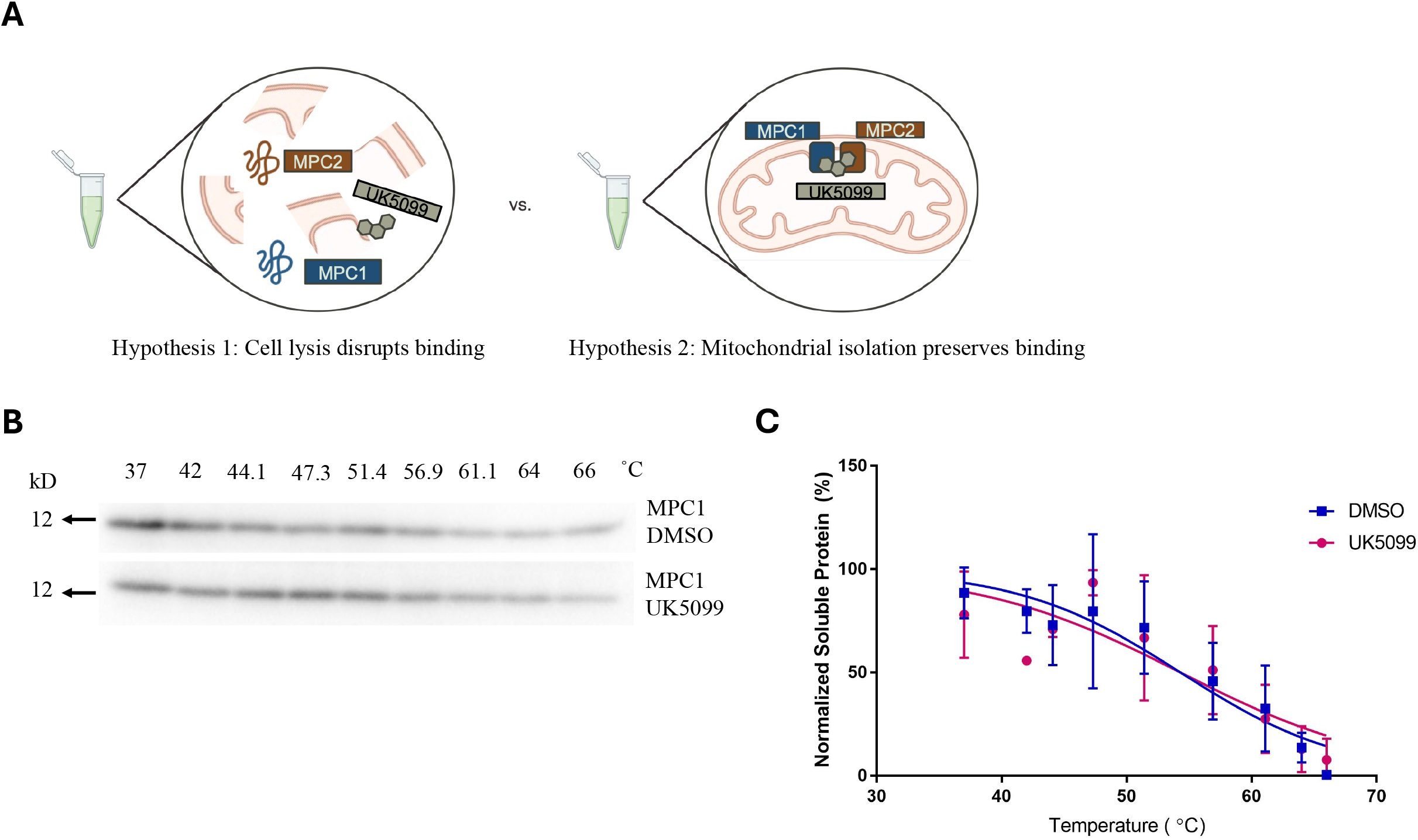
Mitochondria lysis disrupts binding of UK5099 to MPC. (A) Schematic representing hypothesis that cellular lysis disrupts mitochondrial membrane integrity whereas with use of mitochondrial isolation it is preserved. (B) Western blot image of MPC1 in the soluble fraction between the temperatures 37 and 66°C in control versus UK5099 treated samples. (C) The samples were fitted to the Boltzmann Sigmoid curve, and no significant difference was observed in the MPC1 melting temperature between control and UK5099-treated conditions (n = 1-2, N = 3, extra sum of square F test, p = 0.7611). Blot images are representative of means obtained from independent replicates.

## Conclusion

Here we describe the development and validation of a novel subcellular CETSA protocol for evaluating drug-target interactions with membrane-bound proteins and protein complexes. Using the well-characterized interaction between UK5099 and MPC complex, we demonstrated that subcellular CETSA successfully validates the established direct binding of the drug to its known MPC1:MPC2 mitochondrial membrane targets. The demonstrable thermal stabilization of MPC in isolated mitochondria compared to the lack of thermal shift in the context of whole cell or cellular or mitochondrial isolates highlights how a careful choice of biological sample can overcome limitations in membrane protein-target validation.

Mitochondrial and other subcellular membrane proteins remain challenging to validate in a physiologically relevant context, a key limitation in drug discovery [1]. These proteins play key roles in disease and underscore the need for additional validation efforts to position these proteins as drug targets [6]. Expanding subcellular CETSA to additional mitochondrial membrane proteins and other membrane-bound organelles such as lysosomes, whose isolation has been previously described, allows the analysis of targets that have been previously overlooked [11]. Since different membrane proteins and complexes exhibit distinct intrinsic thermal stabilities and aggregation profiles, optimization of temperature range, incubation time, and protein input will be required on a target-by-target basis when applying subcellular CETSA to other membrane-bound proteins. The technical accessibility, preservation of native membrane architecture, and preservation of native protein conformations however position subcellular CETSA as a readily available tool for establishing the therapeutic potential of these proteins.

## Acknowledgements

The authors thank Dr. Gregory Chin for securing the grant that funded this research and for providing valuable feedback on the manuscript.

## Notes

### Competing Interest Statement

The authors have declared no competing interest.

