## Supplemental Figure 1 for "Cellular thermal shift assay of subcellular isolates for evaluating drug-membrane target interactions"

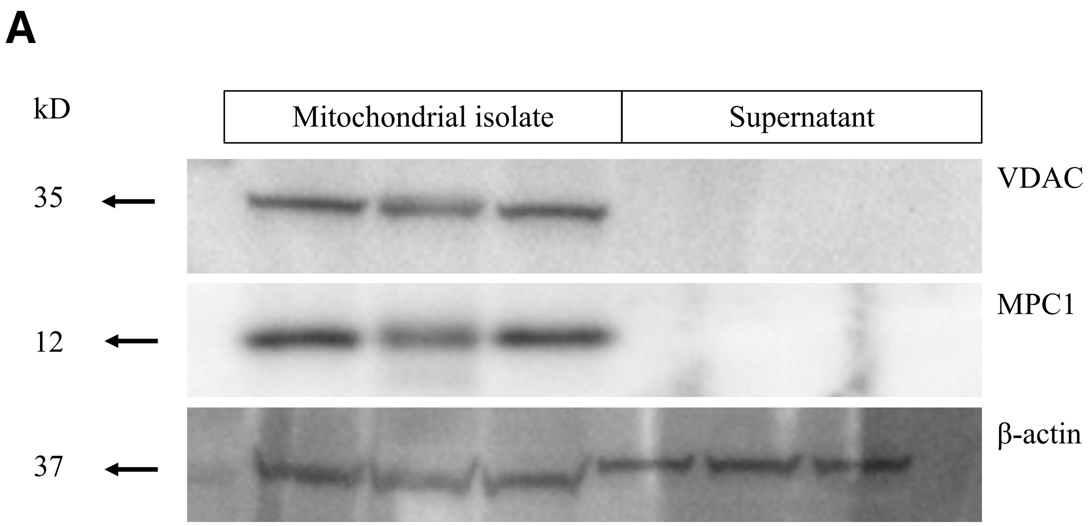


**Supplemental Figure 1: Enrichment of intact mitochondria in the mitochondrial isolate fraction.** Representative Western blot images of VDAC, MPC1, and ß-actin in the mitochondrial pellet and supernatant following the final mitochondrial isolation step. Lack of VDAC and MPC1 in the supernatant is consistent with intact mitochondria specific to the mitochondrial fraction. Blot images are representative of means obtained from independent replicates.
